# Early low-level developmental arsenic exposure impacts mouse hippocampal synaptic function

**DOI:** 10.1101/2021.04.15.440033

**Authors:** Karl F.W. Foley, Daniel Barnett, Deborah A. Cory-Slechta, Houhui Xia

## Abstract

**Background:** Arsenic is a well-established carcinogen known to increase all-cause mortality, but its effects on the central nervous system are less well understood. Recent epidemiological studies suggest that early life exposure to arsenic is associated with learning deficits and behavioral changes, and increased arsenic exposure continues to affect an estimated 200 million individuals worldwide. Previous studies on arsenic exposure and synaptic function have demonstrated a decrease in synaptic transmission and long-term potentiation in adult rodents, but have relied on in vitro or extended exposure in adulthood. Therefore, little is known about the effect of arsenic exposure in development.

**Objective:** Here, we studied the effects of gestational and early developmental arsenic exposure in juvenile mice. Specifically, our objective was to investigate the impact of arsenic exposure on synaptic transmission and plasticity in the hippocampus.

**Methods:** C57BL/6 females were exposed to arsenic (0, 50ppb, 36ppm) in their drinking water two weeks prior to mating and continued to be exposed to arsenic throughout gestation and after parturition. We then performed field recordings in acute hippocampal slices from the juvenile offspring prior to weaning (P17-P23). In this paradigm, the juvenile mice are only exposed to arsenic in utero and via the mother’s milk.

**Results:** High (36ppm) and relatively low (50ppb) arsenic exposure both lead to decreased basal synaptic transmission in the hippocampus of juvenile mice. There was a mild decrease in paired-pulse facilitation in juvenile mice exposed to high, but not low, arsenic, suggesting the alterations in synaptic transmission are primarily post-synaptic. Finally, high developmental arsenic exposure led to a significant increase in long-term potentiation.

**Discussion:** These results suggest that indirect, ecologically-relevant arsenic exposure in early development impacts hippocampal synaptic transmission and plasticity that could underlie learning deficits reported in epidemiological studies.

## Introduction

Early life exposure to toxic chemicals and environmental pollutants is associated with learning deficits and behavioral changes (Cory-Slechta et al., 2018; Liu et al., 2014; Santucci et al., 1994). An estimated 200 million people worldwide are exposed to arsenic concentrations in drinking water that exceed the World Health Organization’s recommended limit, 10 parts per billion (ppb) (Naujokas et al., 2013). Exposure to concerning levels of arsenic is not limited to toxic waste sites. Rather, arsenic levels commonly exceed 10ppb in domestic wells throughout the United States, especially in the southwest. While arsenic levels are kept below 10ppb in municipal water supplies, private wells are unregulated and arsenic levels exceed 10ppb in 20 out of 37 principal aquifers in the United States (DeSimone et al., 2015). Even mild increases in arsenic exposure are of concern, as exposure is associated with numerous adverse health outcomes and increased all-cause mortality (Naujokas et al., 2013).

In January 2006, the maximum contaminant level (MCL) of arsenic in public water systems was lowered from 50ppb to 10ppb, in compliance with a previous United States Environmental Protection Agency (EPA) ruling (2001). This change was enacted due to several epidemiological studies demonstrating increased risk of cancer. While acute, high-level arsenic exposure was known to be associated with peripheral neuropathy, relatively little was known about the neurological consequences of chronic, low-level arsenic exposure (National Research Council (NRC) 1999). However, more recent epidemiological studies have demonstrated that arsenic exposure is associated with deficits in cognitive and motor functions in children and adults (Calderon et al., 2001; O’Bryant et al., 2011; Rosado et al., 2007; Tolins et al., 2014; Tsai et al., 2003; Tyler and Allan, 2014; Wang et al., 2020; Wasserman et al., 2004). Additionally, a recent study suggests inequalities in arsenic exposure reductions following the 2006 change in the MCL, such that there was a higher concentration of arsenic in public water systems serving Hispanic and tribal communities, small rural communities, and Southwestern U.S. communities (Nigra et al., 2020). Given the relatively recent change in the arsenic MCL in public water in the U.S., the continued high exposure in many regions worldwide, and the known vulnerability of the developing brain to toxicants and pollutants underscores the critical need to understand the effects of early life exposure to arsenic.

Electrophysiological studies in arsenic-exposed rodent models have begun to shed light on potential mechanistic underpinnings of the associated cognitive deficits. Rodents exposed to high arsenic concentrations throughout early development and adulthood demonstrate a decrease in synaptic transmission and long-term potentiation (LTP) (Nelson-Mora et al., 2018) in the hippocampus that may be secondary to altered glutamate transport (Siddoway, 2011). Similar changes in synaptic transmission and plasticity have been demonstrated by ex vivo exposure of hippocampal slices to arsenite metabolites (Kruger et al., 2006; Kruger et al., 2007). However, the electrophysiological effects of early developmental arsenic exposure have not been distinguished from chronic adulthood exposure. Because arsenic can cross the placenta and enter a mother’s milk (Concha et al., 1998), we reasoned that gestational and post-parturition arsenic exposure could cause changes in synaptic transmission and plasticity even without adulthood exposure. Further, the effects of continued adulthood exposure on hippocampal synaptic function could differ from the effects of early developmental exposure alone. For example, whereas acute ex vivo exposure to arsenic metabolites attenuates LTP in hippocampal slices from adult rats, it facilitated LTP in young rats (Kruger et al., 2009). Here, we studied the effects of in vivo arsenic exposure during gestation and early development by exposing dams to a high level of arsenic (36ppm) or a low level (50ppb), the MCL for public drinking water in the United States prior to 2006. Of note, in this paradigm, the dam is exposed to arsenic directly via drinking water, whereas the offspring are exposed indirectly via placental transmission and the mother’s milk. Surprisingly, we found that maternal exposure to even low levels of arsenic, i.e., 50ppb, impairs synaptic transmission in the hippocampus of offspring. Additionally, we observed different effects of arsenic exposure in our juvenile mice than what has been previously reported in adult mice (Table 1).

## Materials and Methods

### Arsenic Exposure

All experimental protocols were approved by the Institutional Animal Care and Use Committee of the University of Rochester and carried out in compliance with ARRIVE guidelines. Given that arsenic(V) acid salt (arsenate) is the most common form of arsenic in groundwater (Cullen and Reimer, 1989), we utilized sodium arsenate dibasic heptahydrate (Na_2_HAsO_4_ · 7H_2_O; hereon, arsenic), obtained from MilliporeSigma (A6756). C57BL/6 females were exposed to arsenic (0, 50ppb, 36ppm) in their drinking water (distilled deionized H_2_O) starting at six weeks of age. Breeding began at two months of age and arsenic exposure continued after parturition to simulate protracted human exposure conditions. The juvenile offspring were then used for experiments prior to weaning (P17-P23), such that the pups are still nursing-dependent. Both male and female pups were used for experiments. Mice were maintained with a 12:12 hour light:dark cycle, constant temperature of 23°C and ad libitum feeding. An overview of arsenic exposure is shown in Figure 1. In this paradigm, the juvenile mice are exposed to arsenic in utero and via the mother’s milk. Fresh arsenic solutions were prepared and exchanged every 2-3 days to avoid oxidation.

**Figure 1:**
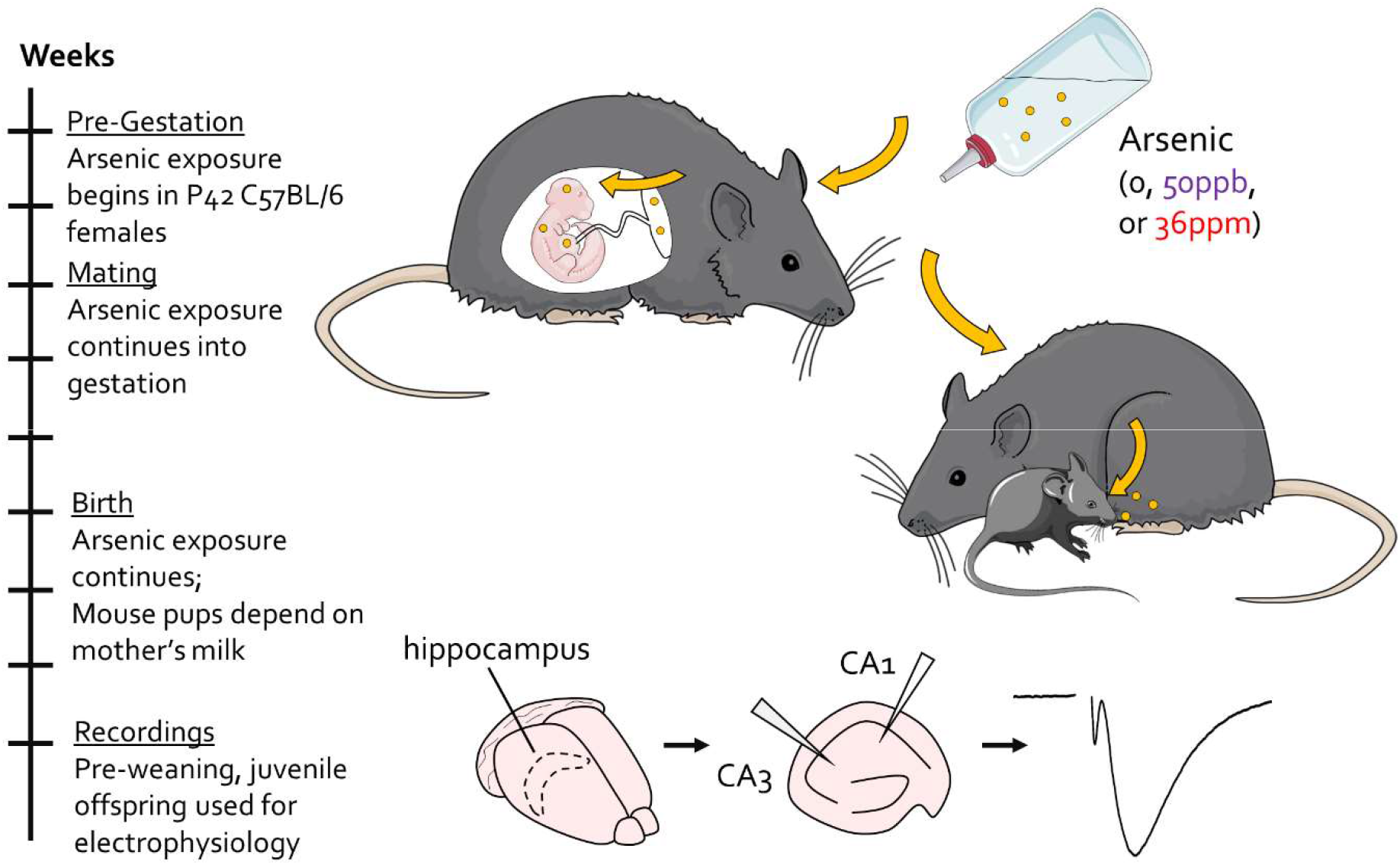
Timeline of arsenic exposure and experiments.

### Electrophysiology

Acute hippocampal slices were prepared from male and female juvenile mice (P17-P23) prior to weaning. 400μm thick hippocampal slices were prepared after decapitation and rapid extraction of the brains into ice-cold artificial cerebrospinal fluid (ACSF). Slices were then allowed to recover in room temperature (RT) ACSF for at least one hour prior to experiments. Field recordings were conducted at Schaffer collateral-CA1 synapses in RT ACSF at a flow rate of 2-3mL/min. A borosilicate recording electrode (1-3MΩ) filled with 1M NaCl was placed in CA1 stratum radiatum and a monopolar stimulating electrode placed on Schaffer collaterals between CA3 and CA1. The ACSF solution consisted of, in mM: 120.0 NaCl; 2.5 KCl; 2.5 CaCl_2_; 1.3 MgSO_4_; 1.0 NaH_2_PO_4_; 26.0 NaHCO_3_; and 11.0 D-glucose. ACSF was aerated with carbogen (95% O_2_, 5% CO_2_) throughout slice preparation, incubation, and recordings. Basal synaptic transmission was assessed by input-output (IO) curves, comparing the fiber volley, a more direct measure of axonal stimulation, to the field excitatory post-synaptic potential (fEPSP) slope. LTP was induced by a single tetanus of one second, 100Hz stimulation. Electrophysiology recordings were collected with a MultiClamp 700A amplifier (Axon Instruments), PCI-6221 data acquisition device (National Instruments), and Igor Pro 7 (Wavemetrics) with a customized software package (Recording Artist, http://github.com/rgerkin/recording-artist).

### Analysis

Electrophysiology data was analyzed using R and GraphPad Prism. To generate IO curves, fiber volley amplitudes were binned at ±0.05 mV, with the exception of 0.025 (0,0.025], 0.05 (0.025,0.05], and 0.1 mV (0.075,1.5]. To measure the magnitude of LTP, we normalized the last five minutes of fEPSPs (25-30 minutes after LTP induction) to the 10 minute baseline. Males and females were pooled for all analyses, with no sub-analysis of the effect of sex. Statistical significance between means was calculated using t-tests or two-way ANOVAs and Dunnett posthoc comparisons, with arsenic exposure and stimulation (fiber volley; inter-pulse interval) as factors.

## Results

Juvenile mice (P17-P23) exposed to either a low (50ppb) or high level (36ppm) of arsenic in utero and in early development exhibit a significant decrease in basal synaptic transmission in the hippocampus at the Schaffer collateral-CA1 synapse (Fig. 2A; two-way ANOVA, arsenic exposure: F(2,216)=22.31, p<0.0001; fiber volley: F(7,216)=99.87, p<0.0001). Interestingly, low and high arsenic exposure levels result in a similar decrease, such that high exposure does not reduce transmission beyond the deficit seen with low-level exposure. However, the two arsenic exposure levels differ in their effects on short-term pre-synaptic plasticity, as assessed by paired-pulse facilitation (PPF) (Fig. 2B). High arsenic exposure reduced PPF at short inter-pulse intervals (both 25ms and 15ms IPI, two-way ANOVA, arsenic exposure: F(2,242)=11.93, p<0.0001; IPI: F(5,242)=49.22, p<0.0001; Dunnett’s post-hoc, p<0.05), whereas there is no significant change from control with low-level arsenic exposure. There was no significant difference between groups at longer inter-pulse intervals.

**Figure 2:**
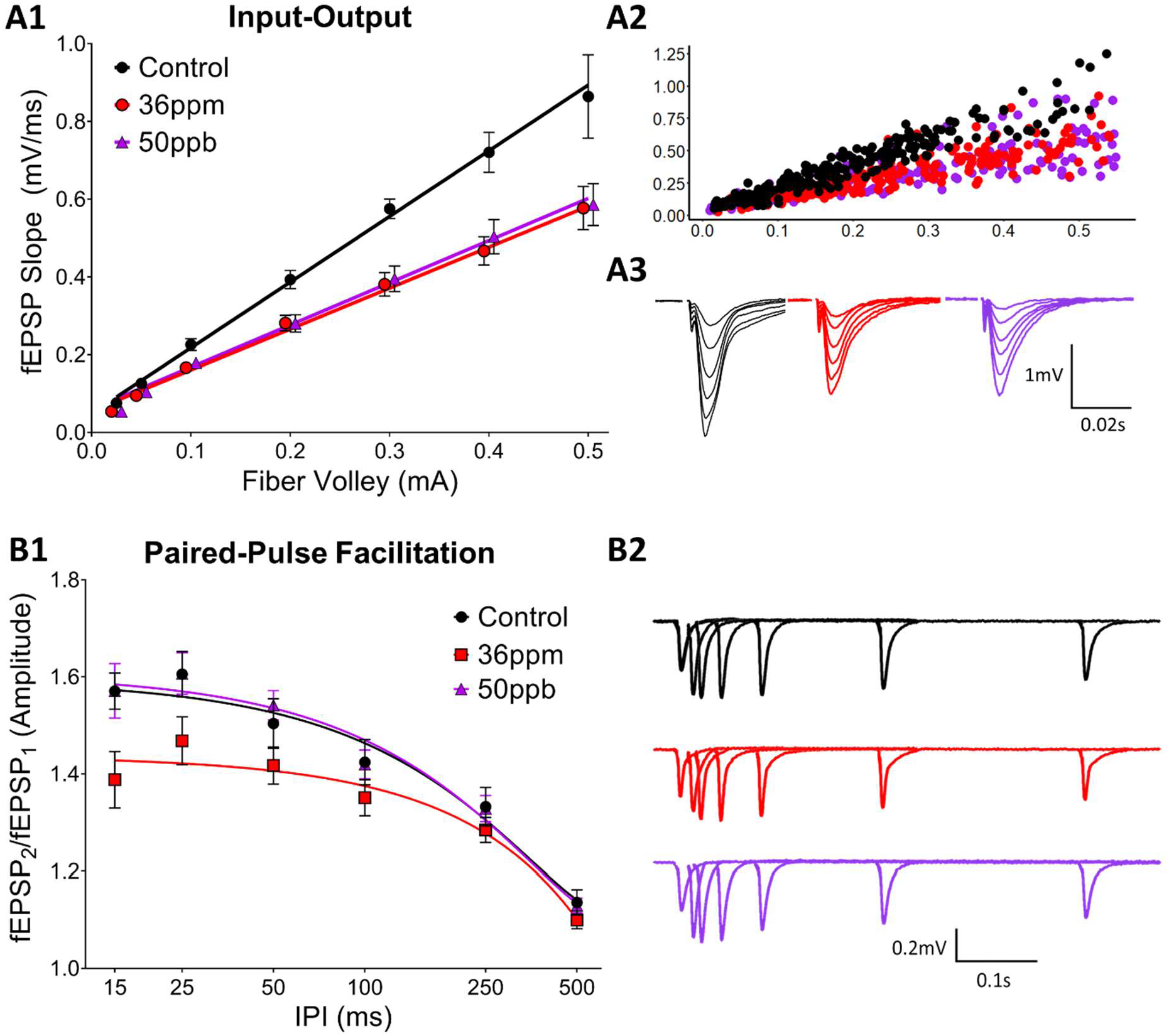
Effect of arsenic on hippocampal basal synaptic transmission (A) and short-term presynaptic plasticity (B), as assessed by input-output curves and paired-pulse facilitation, respectively. The averaged responses from all experiments are shown in A1 and B1 and waveforms from individual experiments in A3 and B2. Pre-binned, individual values from all input-output experiments are shown in A2.

To assess long-term changes in synaptic plasticity, we gave high-frequency stimulation to induce LTP, a cellular model for neural circuit development as well as learning and memory. Surprisingly, high arsenic exposure levels increased LTP by about 11% (Fig. 3; 36ppm: 29%±0.03, n=13; control: 18%±0.04, n=7, p<0.05). Developmental exposure to low-level arsenic led to an 8% increase in LTP compared to control (50ppb: 26%±0.06, n=7), falling between the control and high arsenic exposure, but did not reach statistical significance.

**Figure 3:**
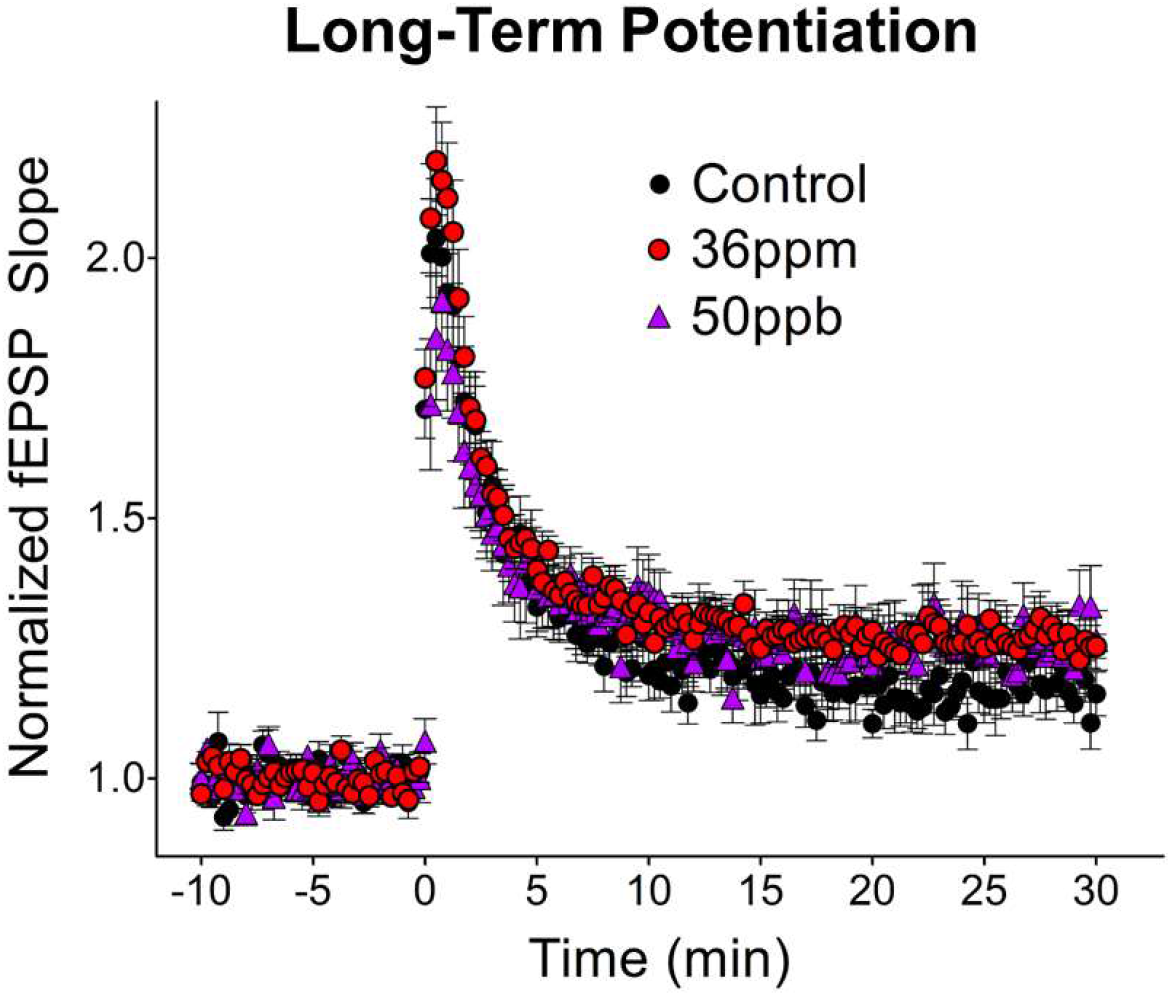
Effect of arsenic on arsenic on hippocampal synaptic plasticity. Long-term potentiation induced by 1×100Hz stimulation.

## Discussion

Our findings show that early developmental arsenic exposure results in significant changes to hippocampal synaptic transmission and plasticity. The most interesting result from our study is that a relatively low dose of arsenic exposure decreases synaptic transmission. This is accompanied by no change in PPF, an indicator of glutamate release in presynaptic neurons; in our case, CA3 neurons. Therefore, our studies suggest that a low dose of arsenic led to a postsynaptic change in CA1 neurons contributing to the decrease in synaptic transmission. This is consistent with the decrease in neurite number and complexity observed in a cell culture model of arsenic exposure (Frankel et al., 2009). On the other hand, a high level of arsenic decreased PPF, suggesting that glutamate release is increased in presynaptic CA3 neurons. However, there is no overall synaptic transmission change between two doses of arsenic (50ppb and 36ppm). This is consistent with the notion that the higher concentration of arsenic caused a progression of changes to the hippocampal circuitry, such that there was a compensatory increase in glutamate release from presynaptic CA3 neurons in response to the changes in postsynaptic CA1 neurons for synaptic transmission.

Our data also indicate that both low and high levels of arsenic led to a trend of increased LTP expression, with an 8 and 11% increase respectively. Our studies build upon and extend previous studies that utilize acute in vitro exposure and chronic in vivo adulthood exposure. First, our findings replicate the decrease in hippocampal basal synaptic transmission observed both in acute, high-concentration in vitro exposure of arsenite metabolites in young rat hippocampal slices (Kruger et al., 2006; Kruger et al., 2007; Kruger et al., 2009) as well as chronic, high-concentration (20ppm) in vivo exposure in adult mice (Nelson-Mora et al., 2018). Second, we found that the decrease in PPF previously observed with high-concentration arsenic exposure in adult mice (Nelson-Mora et al., 2018) was also observed in our high-concentration exposure in juvenile mice (Fig. 2B). Importantly, in our model, changes in basal synaptic transmission were observed even with low-concentration (50ppb) gestational and early developmental exposure. Third, our findings reveal differences between juvenile and adult mice exposed to arsenic. Whereas a previous study found a decrease in the degree of LTP in mice exposed to high-concentration arsenic from gestation through adulthood (Nelson-Mora et al., 2018), we observed an increase in LTP in our juvenile mice. Together, these findings suggest a progression of changes induced by arsenic exposure, where LTP is facilitated in juvenile mice but attenuated with prolonged exposure, consistent with differential effects depending upon the timing of exposure, i.e., the critical window of exposure.

Most strikingly, the pups in our experiments had little direct exposure to arsenic, rather they were exposed in utero and via the mother’s milk, as nursing typically continues into the third week of life. Previous research suggests that the transmission of arsenic through the placenta is higher than transmission into breast milk (Carignan et al., 2015; Concha et al., 1998). Therefore, it is possible that our results are primarily due to the gestational, rather than developmental, exposure in our paradigm. The 50ppb findings are of particular interest for several reasons: 1) 50ppb was the effective arsenic MCL in the United States for decades prior to 2006; 2) the permissible level of arsenic continues to be above 10ppb in many countries; and 3) mean arsenic levels exceed 10ppb in many common beverages and 50ppb in many common foods (Wilson, 2015). The current results suggest that indirect, ecologically-relevant arsenic exposure in early development impacts hippocampal synaptic transmission.

## Acknowledgments

This study was funded by a Pilot Grant from the University of Rochester Environmental Health Sciences Center (P30 ES00247). Figure 1 was created in part using Servier Medical Art images (https://smart.servier.com), licensed under a Creative Commons Attribution 3.0 Unported License.

